# Snaptron: querying and visualizing splicing across tens of thousands of RNA-seq samples

**DOI:** 10.1101/097881

**Authors:** Christopher Wilks, Phani Gaddipati, Abhinav Nellore, Ben Langmead

## Abstract

As more and larger genomics studies appear, there is a growing need for comprehensive and queryable cross-study summaries. Snaptron is a search engine for summarized RNA sequencing data with a query planner that leverages R-tree, B-tree and inverted indexing strategies to rapidly execute queries over 146 million exon-exon splice junctions from over 70,000 human RNA-seq samples. Queries can be tailored by constraining which junctions and samples to consider. Snaptron can also rank and score junctions according to tissue specificity or other criteria. Further, Snaptron can rank and score samples according to the relative frequency of different splicing patterns. We outline biological questions that can be explored with Snaptron queries, including a study of novel exons in annotated genes, of exonization of repetitive element loci, and of a recently discovered alternative transcription start site for the ALK gene. Web app and documentation are at http://snaptron.cs.jhu.edu. Source code is at https://github.com/ChristopherWilks/snaptron under the MIT license.

## 1 Introduction

The Sequence Read Archive (SRA) is a huge and valuable repository of public and controlled-access sequencing data, spanning over 9 petabases and doubling in size every 18-20 months. Such archives allow researchers to reproduce past studies, combine data in new ways, and access unique datasets that would otherwise be too expensive or difficult to obtain. But researchers struggle to take full advantage of archived data. There is no convenient way to pose scientific questions against the archives without first downloading and reanalyzing data. The situation is analogous to the early days of the World Wide Web, when content was accessed at well known addresses via transport protocols (FTP, HTTP). The web became vastly easier to use with the advent of search engines: crawlers, indexes, and ranking algorithms made it possible for users to filter the web for content relevant to their queries.

Snaptron is a search engine that enables study of splicing patterns in large, pre-analyzed collections of human RNA sequencing (RNA-seq) samples. Snaptron answers queries via a Representational State Transfer (RESTful) web service interface. Driving Snaptron is a query planner that combines the strengths of different indexing strategies (R-trees, B-trees and term-document inverted indices) to rapidly address user queries. The data used to service a given query can be a mix of genomic interval data, numeric values associated with genomic intervals, and free- or controlled-text from associated metadata. The REST interface can be queried via HTTP with no software installation necessary. Alternately, Snaptron can be queried via a client script that provides a richer set of queries. Users may also download the (large) files used to populate Snaptron’s database as well as the Snaptron server software to create a fully local Snaptron installation.

While past efforts address the problem of enabling efficient cross-study queries, most have focused on genotype rather than expression or splicing data. GEMINI [1] and Genome Query Tools (GQT) [2] are complementary tools for indexing and querying genotypes from many individuals, facilitating computation of genomewide summaries over subsets of individuals. BGT [3] builds on the positional Burrows-Wheeler Transform (PBWT) [4] to provide similar functionality, including region-specific queries. The ExAC [5] browser and REST allow querying of frequencies of genetic variants, as summarized over 90,000 re-analyzed exomes.

Other past efforts sought to enable querying of expression data in particular. Solomon and Kingsford propose Sequence Bloom Trees (SBTs) [6] for indexing the raw reads from many sequencing samples. They indexed 2,652 human RNA-seq experiments and queried the index using known transcript sequences. The index reports which samples contain the query string, but it is up to the user to build the index and to reduce a biological question into such a presence/absence query. The Expression Atlas [7] summarizes a curated subset of the data in ArrayExpress [8] and enables querying of baseline gene expression and differential gene expression across groups in certain studies. But it enables only gene-level queries, and only performs differential gene expression comparisons over manually-selected subsets of the samples in the archive.

Another focus of past work has been on computational problems that arise when working with many genomic intervals. BEDTools [9] and GenomicRanges [10] are widely used tools for working with intervals. The Genome Query Language (GQL) [11] is a SQL-like language for searching and joining aligned reads with other genome-mapped data. GORPipe [12] provides an interface to genomic interval based data, where queries are accelerated via file seeks on tabular genomic data files sorted by start-positions.

Our goal is to create a full suite of search-engine software for summarizing expression and splicing data. The project began with a “crawling” effort, described previously [13,14], where we used the Rail-RNA software tool to analyze tens of thousands of RNA-seq samples in a uniform fashion. Here we present the Snaptron tool, which can rapidly answer sophisticated queries with respect to splicing and expression data as well as sample metadata. Snaptron makes it easier to leverage large amounts of public data in data-to-day research, without needing users to be overly concerned with lower-level computational ideas (indexing schemes, data structures) used in the system itself. Snaptron is particularly useful for lending additional context and support to hypotheses related to splicing.

In Methods we describe the design of Snaptron and results from the performance profiling that inform the current design. In Results, we describe analyses leveraging both the REST interface and the command-line client for Snaptron. These are examples of analyses that users can perform using Snaptron queries. The Snaptron REST service and documentation are available at: http://snaptron.cs.jhu.edu. The Snaptron software is freely available under an MIT license from: https://github.com/ChristopherWilks/snaptron. Software for performing the analyses described here are available as described in Supplementary Note 1.1.

**Figure 1:**
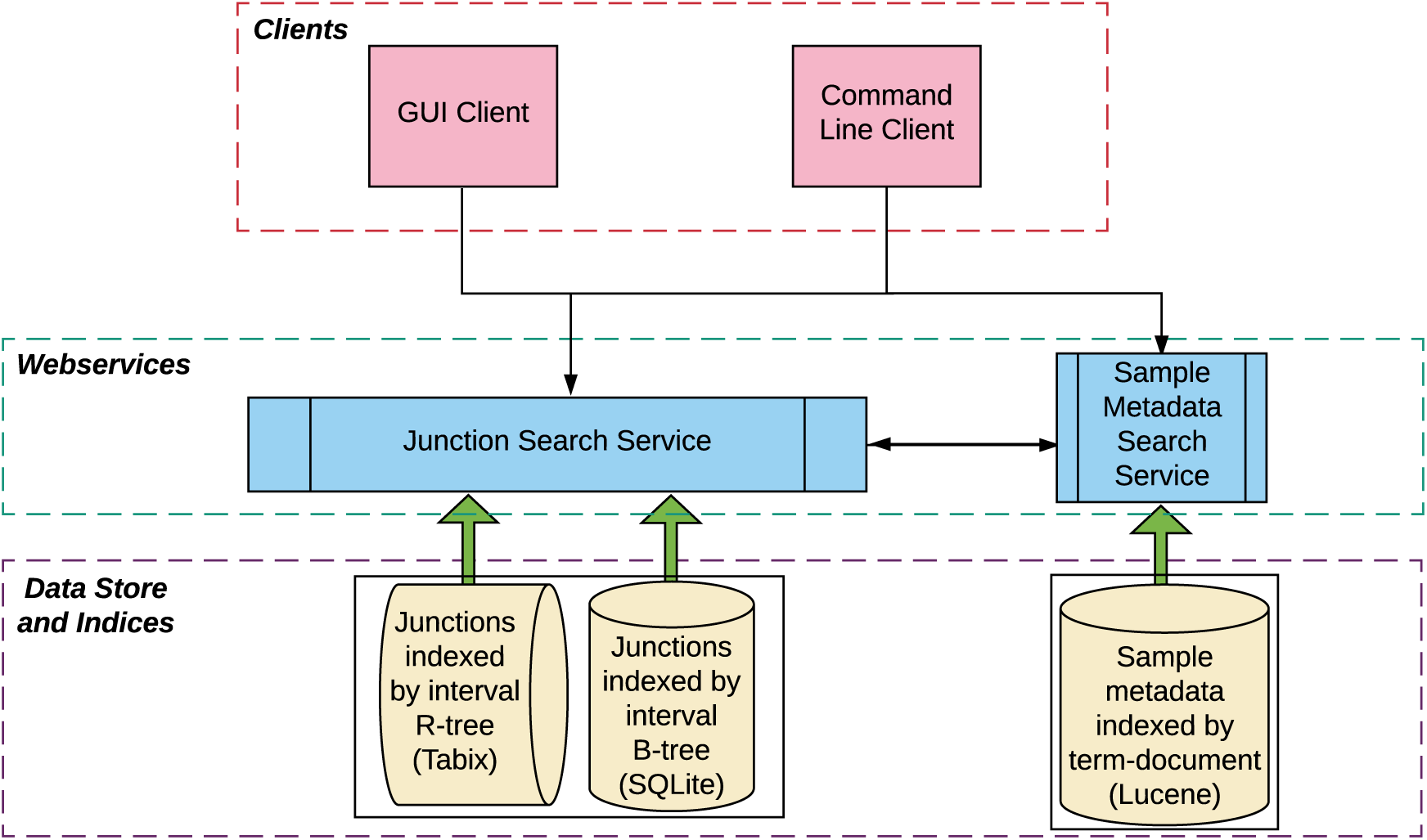
The Snaptron architecture consists of three layers (from the bottom up) including data and associated indices (Tabix, SQLite, and Lucene), webservices and processing (Python), and finally the clients (NodeJS and Python). Queries flow down from the clients to and between the web services (black arrows) while data flows back from the indices through the webservices to the clients (large, green arrows).

## 2 Methods

### Crawling and summarizing

To produce the splicing data served by Snaptron, we used Rail-RNA [13] to analyze many archived human RNA-seq samples. Rail-RNA is a scalable spliced aligner designed to analyze many samples at once. Among Rail-RNA's outputs is a table summarizing evidence for exon-exon splice junctions across all samples.

Each row describes a junction, its strand and coordinates, and the number of reads spanning the junction for each sample where it appears. We also created tables detailing metadata for each sample. This is the source material for Snaptron as well as for the `intropolis` resource [14]. `Intropolis` makes junction data available for bulk download but without an indexing facility and without an interface for querying the data. Further details on alignment of GTEx samples are contained in [15], while details on alignment of other SRA samples as well as TCGA samples are contained in [16].

Snaptron further adds auxiliary information to each junction:

- Gene annotation status (discussed below)
- Count of samples with one or more reads covering the junction
- Sum, average, and median of the junction coverage across samples where the junction occurred at least once

Snaptron allows the user to query any of these four compilations of human RNA-seq samples:

- SRAv1: 43M junctions from 21,504 public samples from the SRA
- SRAv2: 81M junctions from 44,427 public samples from the SRA
- GTEx: 29M junctions from 9,662 samples from the v6 data freeze
- TCGA: 37M junctions from 11,284 samples from TCGA

SRAv1 uses the GRCh37 human reference genome and its coordinate system. SRAv2, GTEx and TCGA use GRCh38 and its coordinates. While raw GTEx and TCGA data are dbGaP-protected, Snaptron’s junction-level summaries are, like the SRAv1 and SRAv2 compilations, publicly accessible. Snaptron indexes each compilation separately.

We used a composite of several gene annotation sources (Supplementary Table 1) to determine annotation status of each junction, and of each junction’s donor and acceptor splice sites. If the junction as a whole appears in an annotation, we mark the junction and its splice sites as annotated. If the junction does not appear as a whole, each splice site is marked according to whether it participates in an annotated splice junction. We used UCSC's liftOver tool to convert annotation sources between genomic coordinate systems.

**Figure 2:**
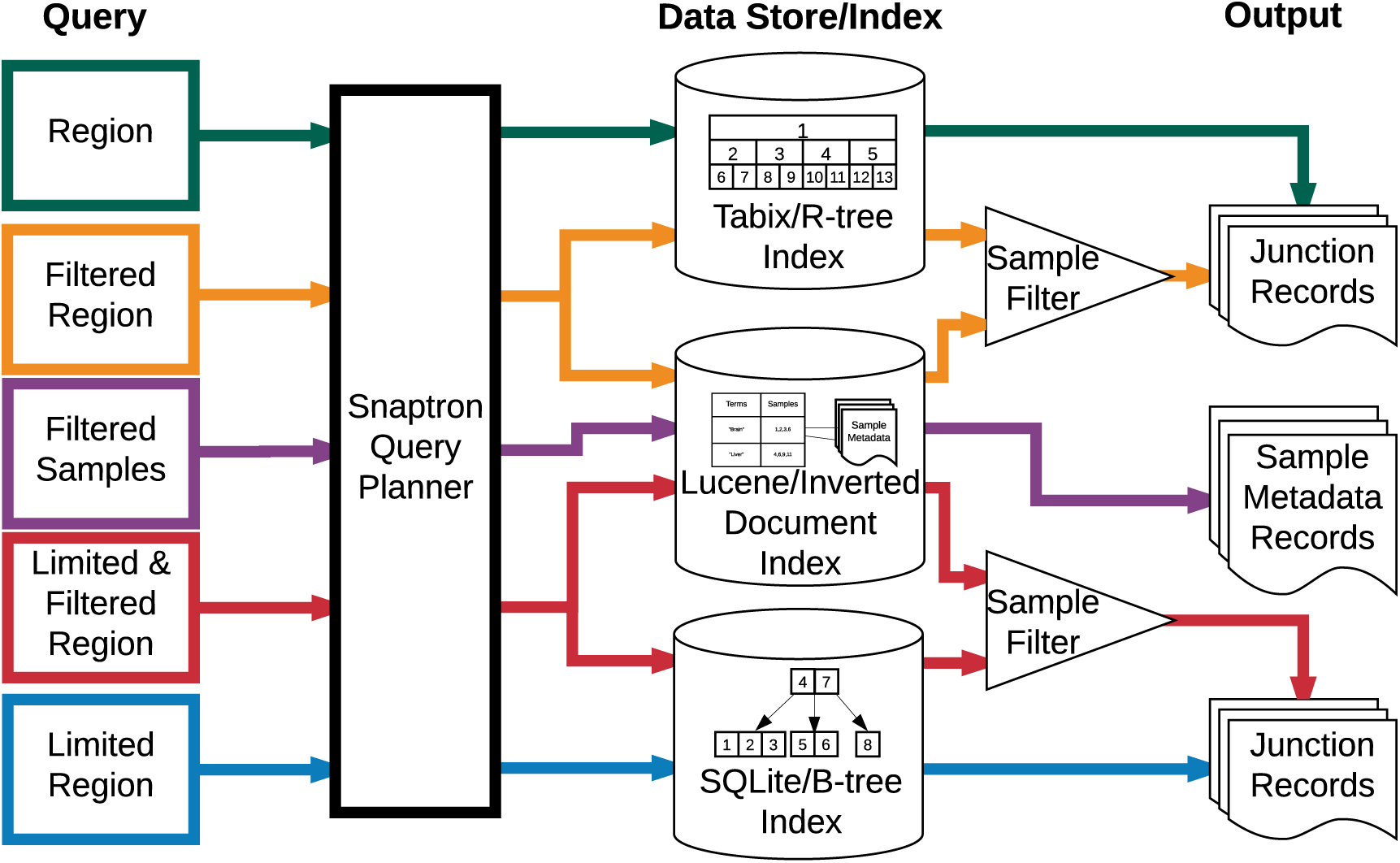
The flow of each query through Snaptron and the type of output it produces. Colors correspond to those used in Table 1.

### Data types

Snaptron uses a hybrid indexing approach that enables efficient querying and retrieval. Queries can be concerned with these distinct but related data types:

- *Genomic intervals,* each consisting of a chromosome and beginning and ending offsets. An interval might represent an exon-exon splice junction or an exon as it appears in a gene annotation. A collection of intervals might represent all exon-exon splice junctions in a sample. Intervals within a collection might overlap, i.e. cover some of the same genomic positions.
- *Integer and floating-point numbers.* For example, Snaptron uses non-negative integers to encode the number of reads spanning an exon-exon junction in a sample. Snaptron also stores pre-calculated summaries such as the average coverage of a junction across all samples it appears in, or the number of samples in which a junction has non-zero coverage.
- *Variable length strings of text*, for sample metadata. Metadata is stored in a combination of semi- and un-structured fields where semi-structured fields here are controlled vocabulary and ID/accession fields. Unstructured data are free-text fields ranging from fragments to full paragraphs. A semi-structured text field might contain metadata of a category that is particularly clearly annotated: i.e. sex and tissue type of a sample from the GTEx project. An un-structured field might contain a study's abstract or a description of how the sample was prepared for sequencing.

The particular query determines which index or combination of indices Snaptron uses to compose the response.

### Region query

A Region (R) query (Table 1) retrieves junction data situated in a given genomic interval. It is handled using Tabix [17] and its associated R-tree index. Such a query might ask for a list of exon-exon junctions that occur in any sample and that overlap a specified genomic interval. An R-tree index is a tree of nested multi-dimensional bounding rectangles. An R-tree node corresponds to the minimum bounding rectangle for the points below in the tree. Since we are working with one-dimensional intervals, the bounding rectangles are simply line segments, and the R-tree is essentially an interval tree [17] [18]. A junction and all associated data is stored in stored the lowest R-tree node fully containing the spliced interval. Data associated with a junction includes its strand, splice motif, annotation status, and its depth of coverage in each sample.

**Table 1:**
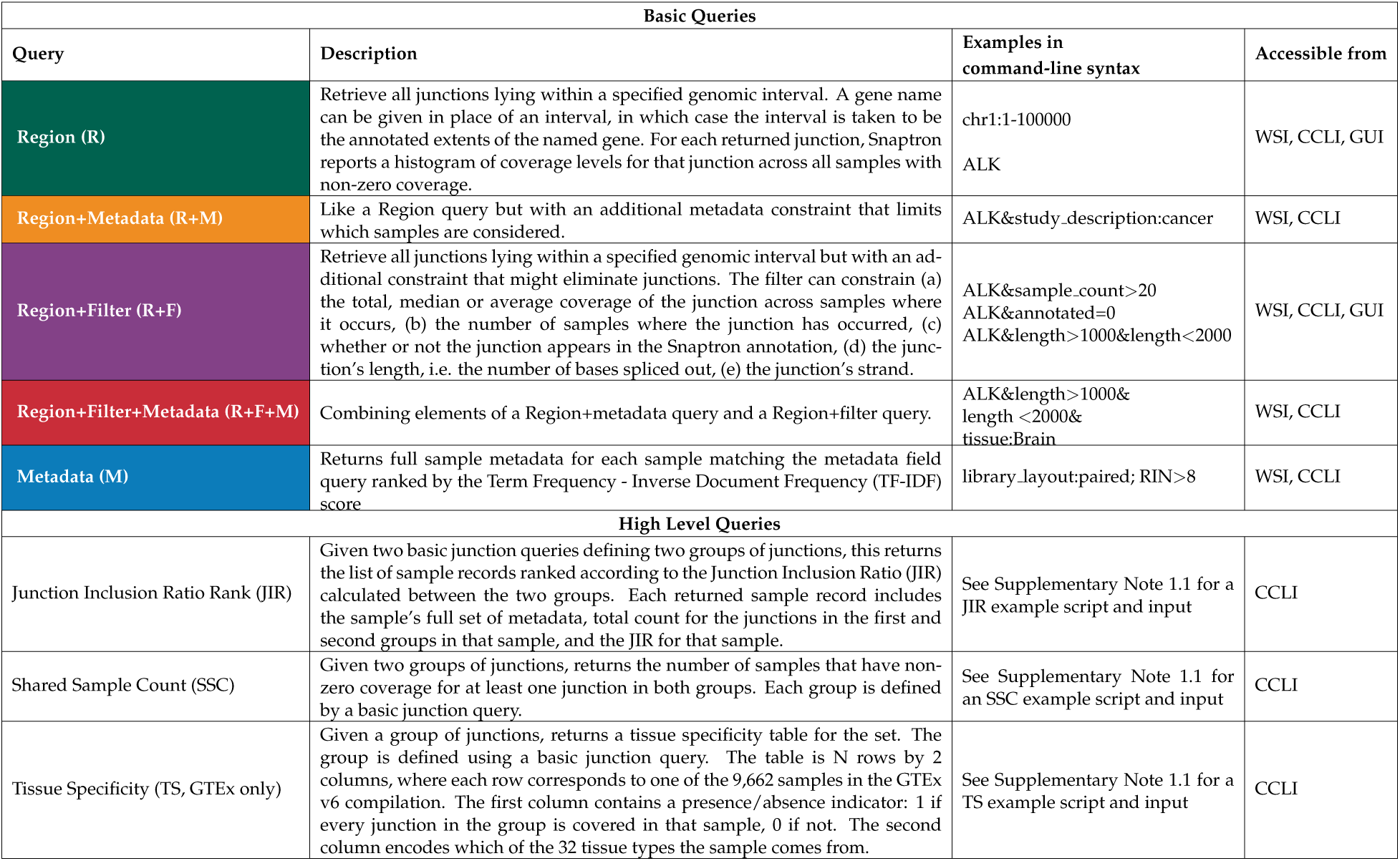
Description of basic and high-level queries supported by Snaptron.

Querying a human-scale Tabix index on the current Snaptron server takes a few seconds, including overhead added by Python and Snaptron (Figure 3). However, the performance of the Tabix R-tree depends on whether the indexed dataset is compressed. We compared the relative performance of the SQLite B-tree and the Tabix R-tree, using Tabix in both compressed and uncompressed modes, and found that uncompressed Tabix outperforms SQLite but SQLite outperforms compressed Tabix (Figure 3). Thus, Snaptron uses the Tabix R-tree in uncompressed mode. Further discussion of these experiments is in Supplementary Note 1.2.

**Figure 3:**
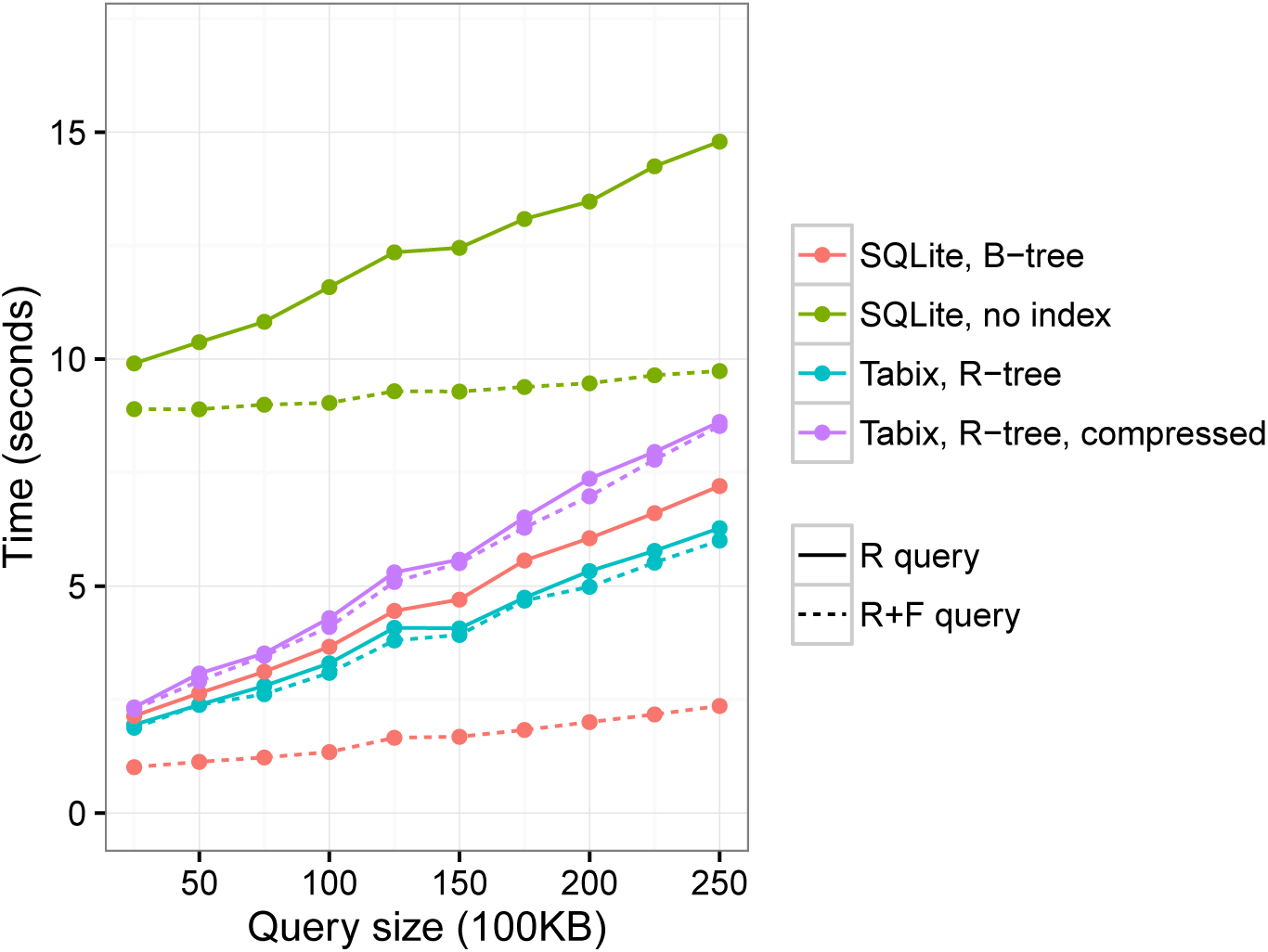
Snaptron query wall-clock times for R and R+F queries of increasing size. The queries ask for all (for R) or some (for R+F) junctions overlapping an increasingly large prefix of chromosome 1. The region grows from a 2.5M-base prefix (leftmost) to 25M bases (rightmost) in 2.5M increments. The R+F constraint additionally requires all junctions returned to have samples_count>=100. The number of junctions returned by the R query range from 350K for the smallest (leftmost) to 1.5M for the largest (rightmost). The number of junctions returned by the R+F query range from 7.3K for the smallest to 28K for the largest. All data was uncompressed except where noted. Details of these runs are provided in Supplementary Note 1.2.

**Figure 4:**
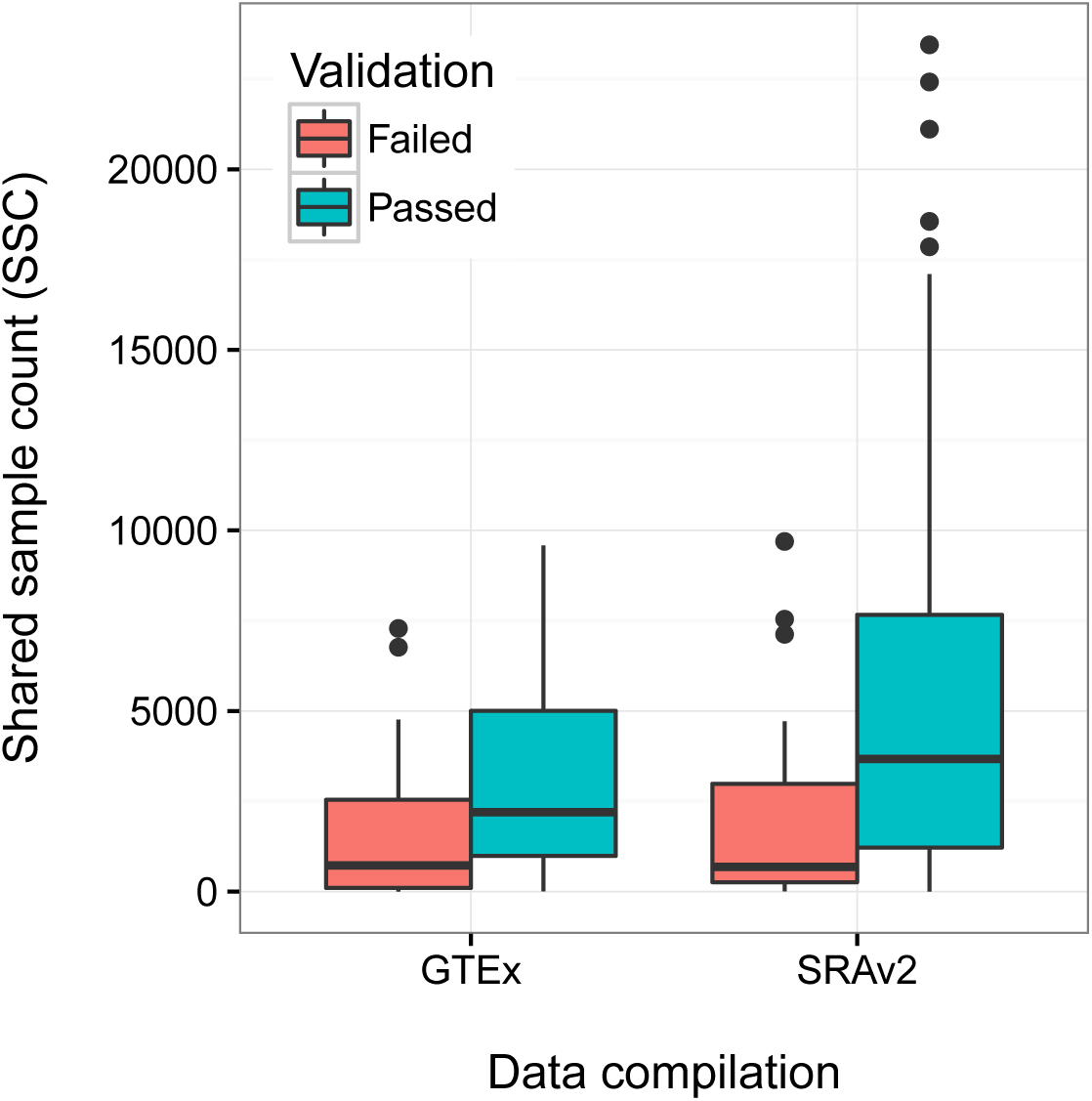
Co-occurring sample counts distinguishing validated from non-validating alternatively spliced exons. For GTEx, Wilcoxon rank-sum *p =* 2e-04. For SRAv2, Wilcoxon rank-sum *p =* 1e-05.

When querying to find junctions that “match” a specified interval, Snaptron allows the user to specify precisely what it means to “match”:

- **contains**: match any junction that falls entirely within the specified interval
- **exact**: match any junction with exactly the specified chromosome, start and end coordinates
- **either**: match when either the start or end coordinate falls inside the interval

### Filtering attributes

A Region + Filter (R+F) query additionally constrains junction attributes. These attributes describe, for example, annotation status, strand, or prevalence. Examples of attribute constraints are given in Table 1. Snaptron uses SQLite and its B-tree index for these queries [19]. Probes into the B-tree index are efficient — logarithmic in the size of the tree — and the tree data is organized in a blocked fashion that enables efficient transfers to and from disk. In a Snaptron B-tree, a key represents a junction, including its chromosome, start coordinate and end coordinate. The full record, including the junction and all associated data (strand, annotation status, coverage in each sample, etc) is stored in a related tree structure.

Some junctions in Snaptron’s compilations are false positives, due to alignment error and other factors [13]. Consequently, we expect a popular query type will be R+F queries requiring the returned junctions to meet a minimum level of prevalence. For instance, a user might ask for “only junctions with coverage ≥ 50” or “only junctions with non-zero coverage in ≥ 1,000 samples.”

While Snaptron could re-use the Tabix R-tree for both R and R+F queries, we found SQLite's B-tree was faster for the R+F case (Figure 3). This is because the junction attribute filter is not handled by Tabix; rather, the F constraint must be handled separately by Snaptron, which parses Tabix output and suppresses records not satisfying the constraint. This adds overhead compared to SQLite, which can naturally combine the interval and attribute constraints in a single action.

### Constraining metadata

Metadata constraints narrow Snaptron’s focus to only those samples with metadata matching or containing key phrases. If we think of the junction evidence as forming a matrix with rows corresponding to junctions and columns to samples, metadata constraints narrow the query's focus to a subset of the columns. A metadata constraint can be used on its own in a Metadata (M) query, or combined with a Region (R) query or Region + Filter (R+F) query. We call the latter two combinations R+M and R+F+M queries (Table 1).

Snaptron uses the popular Lucene [20] inverted indexing system to handle metadata constraints. The Snaptron server includes Lucene indices for each data compilation: SRAv1, SRAv2, GTEx v6 and TCGA. Each index associates over fifty metadata fields with each sample. The exact fields and field contents depend on the data source. Some fields contain unstructured (“free”) text and describe, for example, how the sample was prepared and sequenced or what was being studied. Other fields are semi-structured, using text labels to describe categorical variables, such as whether the reads are paired-end or the sample’s tissue type. For example, the GTEx compilation includes a controlled-vocabulary field describing the tissue of origin, but the SRA compilations do not; that information can often be gleaned from other free text-fields, though sometimes with difficulty [21]. The Lucene index allows searching for key phrases in any metadata field.

### Query planning

Snaptron's query planner determines what index probes are needed and how to combine their output to answer the overall query. Region (R), Region + Filtered (R+F), and Metadata (M) queries are each answered from a different index; R queries use the Tabix R-tree, R+F queries use the SQLite B-tree, and M queries use the Lucene inverted index (Figure 1).

The situation is more complex when a query combines a region constraint with a metadata constraint, as in the case of R+M and R+F+M queries. If we again think of the junction evidence as a matrix with junctions as rows and samples as columns, queries combining R and M constraints are concerned with a subset of columns (M constraint) and a subset of rows (R constraint). Such a query might ask for all junctions in the KCNIP4 gene that appear in at least 10 brain samples.

The problem of handling this query can be decomposed into a few tasks. A “column projection” task determines which samples satisfy the metadata constraint. If C denotes the full set of columns (samples), let *C*′ ⊂ *C* be the subset satisfying the constraint. A “row projection” task determines which junctions (rows) satisfy the region constraint. If *R* denotes the full set of rows (junctions), let *R*′ ⊂ *R* be the satisfying subset. Once *C*′ and *R*′ are known, the final task (“submatrix filtration”) is to determine the subset of *R*′ satisfying the “at least 10” constraint. Submatrix filtration is concerned only with the *R*′ × *C*′ submatrix, so summaries calculated over entire rows or columns from the original matrix cannot be reused here.

The column projection is handled by querying the Lucene index and the row projection is handled by querying the Tabix index. The set of sample IDs returned by Lucene is then converted into an Aho-Corasick automaton. The automaton performs setwise pattern matching on Snaptron’s internal string representation of the matrix rows. Specifically, Snaptron stores a row as a comma-delimited string, with each field containing the concatenation of the sample ID and the read coverage of the junction in that sample. To save space, samples with 0 coverage are not included as fields. The automaton analyzes a single row by consuming the row string’s characters one-by-one and signaling when it has encountered one of the selected columns by entering a special “match” state. Each such match contributes a non-zero entry to the *R*′ × *C*′ submatrix. Once the submatrix is formed, Snaptron re-calculates row-wise summaries (e.g. sum, average, median). Finally, if an attribute filter (F) was specified, it is evaluated with respect to the recalculated summaries to further narrow the list of returned junctions, completing submatrix filtration.

### Higher-level functions

Snaptron supports three higher-level queries showcased in the Results section (Table 1). They are called higher-level because each involves particular sets of junctions defined using “basic” sub-queries.

The *shared sample count* (*SSC*) high-level query returns the number of distinct samples in a compilation that have evidence supporting the co-occurrence of two junctions. Later we show how to use SSC to study the prevalence of putative novel exons. The output of the query can easily be loaded into an R or Python session and analyzed or visualized to better understand the prevalence of a splicing pattern.

The *tissue specificity* (*TS*) high-level query uses the GTEx v6 compilation. The user specifies one or more groups of junctions using one or more region sub-queries, one sub-query per group. The TS query returns an *N* × 2 table, where the *N* rows correspond to all 9.6K samples from the GTEx project and the two columns correspond to (a) whether a junction from every group occurred in that sample, and (b) which of the 32 GTEx v6 tissue types the sample was derived from. This list can then be loaded into Python or R to assess tissue specificity.

The junction-inclusion ratio (JIR) high-level query scores each sample according to a particular overrepresented splicing pattern relative to another. The user specifies two groups of junctions using two region sub-queries. Call these groups A and B. The query calculates the normalized difference between coverage counts of the two groups across all samples containing the junctions. This is the “junction inclusion ratio” (JIR) suggested previously by Nellore et al [14], but with one added to the denominator:

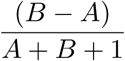

A and B represent the total coverage for the two groups of junctions in the sample. Ranking samples according to JIR reveals the degree to which a splicing pattern is specific to a particular kind of sample.

### Interfaces

Snaptron provides the following interfaces:

- RESTful web service interface (WSI): handles query requests made by a user or by other Snaptron interfaces via HTTP 1.1. Results come in the form of lists of junctions and associated junction data, or lists of samples and associated sample metadata. Queries usually return within seconds.
- Client command-line interface (CCLI): handles both basic queries and high-level queries. High-level queries are decomposed and handled via one or more WSI queries.
- Web-based graphical user interface (GUI): allows a user to visualize splicing patterns across many samples for a given window of the genome. Supports basic queries for populating the window. Queries are decomposed and ultimately handled by the WSI.
- Complete server installation (Local): users can download the underlying Snaptron data and software, build local indices and compilations, and run a local Snaptron service for handling WSI queries. This is for advanced users who require rapid processing of high query volumes.

Users experienced with command-line tools may prefer the direct WSI interface, which has minimal software requirements and responds within seconds in most cases. Users willing to install the lightweight Python CCLI additionally gain access to high-level queries. The CCLI is easily called from wrapper scripts to compose more complex analyses, as shown in our example scripts.

The GUI is ideal for users concerned with a particular gene or genomic region. The GUI will display junctions in that region, distinguishing clearly between annotated and unannotated junctions and using colors to differentiate junctions that occur more or less widely in the dataset. After exploring in the GUI, users can switch to the CCLI or WSI to answer specific questions.

## 3 Results

We describe three analyses that use Snaptron. Each leverages public data to provide context or support for a hypothesis about splicing. Scripts for performing these analyses are available as described in Supplementary Note 1.1. Figure 5 shows screen captures from the Snaptron GUI for each analysis.

**Figure 5:**
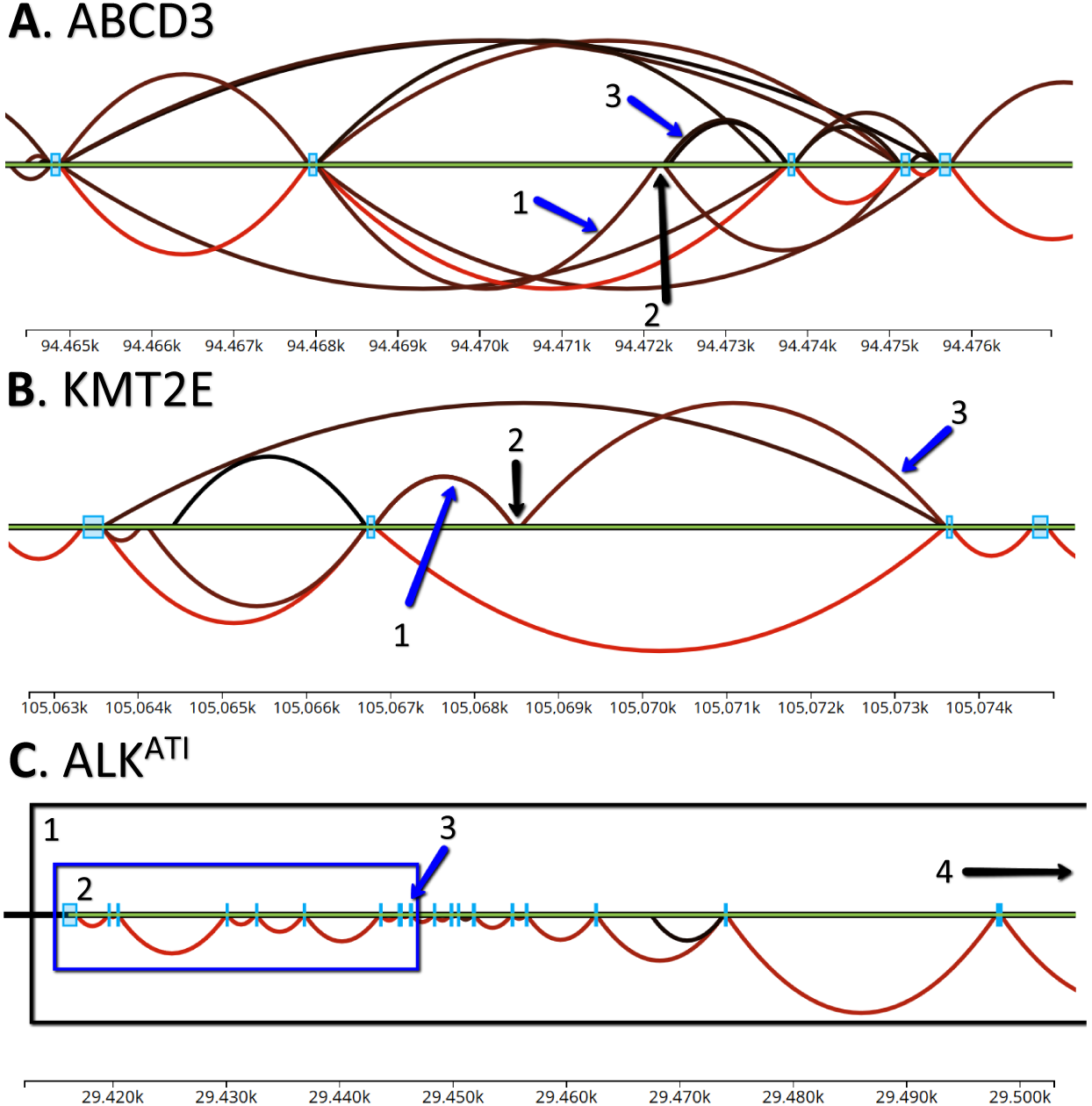
Three Snaptron GUI screen captures corresponding to the three analyses. Green horizontal lines indicate the genome. Arcs indicate exon-exon splice junctions. Colors indicate the number of samples having evidence for the junction, ranging from black (least support) to red (most). Annotated junctions are represented by arcs above the green line, and unannotated junctions by arcs below the line. Light blue rectangles are annotated exons. A) Splice junctions matching the Goldstein et al prediction of a novel alternative exon in the ABCD3 gene. A1 is the 5’ junction, A2 is the novel exon, and A3 is the 3’ junction; B) KMT2E gene and unannotated junctions supporting a REL exonization event. B1 is the 5’ junction, B2 is the REL exon, and B3 is the 3’ junction; C) ALK spliceforms. C1 indicates the full length ALK transcript, C2 is the truncated ALK^ATI^ transcript incorporating only the last 10 exons (ALK is on the reverse strand, and so is laid out right-to-left), C3 is the alternative transcription initiation exon, and C4 is the upstream full transcription initiation site.

### 3.1 Novel Exon Discovery and Evaluation

Snaptron can measure the prevalence of candidate junctions and exons in public RNA-seq data. Candidate junctions need not be annotated; since Snaptron’s junction calls were made without the influence of a gene annotation, it can give support for both annotated and unannotated splicing patterns. We demonstrate this by following the work of Goldstein et al [22], who searched for novel (unannotated) cassette exons in Illumina Human Body Map 2.0 RNA-seq data from 16 normal tissues. A cassette exon was called novel if neither edge coincided with an annotated junction, but the entire exon was located within an annotated gene. Goldstein et al discovered 249 novel exons and validated 216 in a separate cohort using additional paired-end RNA-seq sequencing.

To find evidence for these 249 exons, we posed a high-level Snaptron query that (a) gathered evidence for the exons in the SRAv2 and GTEx compilations, and (b) scored the exons according to shared sample count (SSC), the number of samples with evidence for the exon. The query also constrained the reported junction’s strand to match the strand of the enclosing annotated gene. Further, the query included the “either” modifier to ensure one end of the queried junctions would match exactly with the flanking coordinate of either the 5′ or 3′ end of the predicted exon.

We found that out of 249 putative exons, 236 (94.8%) occurred in both the SRAv2 and GTEx compilations. Of the 236, 204 (86.4%) were among the exons that Goldstein et al validated in a separate cohort, while the remaining 32 were among the exons that failed to validate. We further used the shared sample count (SSC) to score each of the 236 exons and found that the exons that validated by Goldstein et al had significantly higher SSC than those that failed 4. This was true regardless of whether we used the SRAv2 or the GTEx compilation to calculate the score.

This elaborates Goldstein et al's analysis in two key ways. First, Snaptron used public data to score candidate novel exons according to the amount of supporting evidence across tens of thousands of public RNA-seq samples. The scores are valuable both for understanding the degree to which the exons should be considered “novel,” and for prioritizing follow-ups such as PCR experiments. Second, Snaptron’s comprehensive annotation shed further light on the annotation status of the exons. While Goldstein et al determined that the 249 putative novel exons were unannotated at the time, we found 132 were fully annotated by one or more annotation sources used by Snaptron, with the SIBgenes [23] and ACEview [24] tracks annotating most of the 132.

### 3.2 Exonization of Repetitive Elements

Snaptron can also use public data to measure the tissue specificity of a splicing pattern. A repetitive element locus (REL) exonizaton is an instance where a stretch of repetitive sequence (e.g. a SINE or LINE) is spliced into a surrounding gene as an exon. A study by Darby et al [25] reported numerous REL exonization events in human protein-coding genes, including events specific to brain or blood. We used Snaptron to study these events and to measure tissue specificity with respect to the much larger SRAv2 and GTEx compilations.

We began by obtaining coordinates for 5 PCR-validated REL exonization events in three genes (KCNIP4, KMT2E and GLRB) and using the shared sample count (SSC) high-level query to measure the prevalence of the events in the SRAv2 and GTEx collections. We noted that the samples studied by Darby et al, derived from the Stanley brain collection, were not present in these compilations. For both the SRAv2 and GTEx compilations, we found that all five events had a shared sample count of 39 or greater. We also found that none of the junctions flanking the events were fully annotated, in agreement with Darby et al.

We also analyzed the tissue specificity of the 5 REL exons using Snaptron's high-level Tissue Specificity (TS) query (Table 1). We then performed a Kruskal-Wallis rank sum test on the output from the TS query, using the presence/absence results as the data vector and the tissue-annotation results as the group vector. All rank sum tests yielded *P* < 1 · 10^−9^, indicating strong tissue specificity. For example, the REL exon we refer to as `GLRB_1` is present in 33% of the 1,409 samples labeled “Brain” but only 3% of other samples. Similarly, the REL exon `KMT2E_1` is present in 56% of the 102 samples labeled “Bone Marrow” but only 12% of other samples.

### 3.3 ALK and Junction Inclusion Ratio

Snaptron can also be used to study splicing patterns involving many junctions. To demonstrate this, we performed an experiment modeled on Nellore et al’s analysis of the anaplastic lymphoma kinase (ALK) gene’s ALK^ATI^ variant isoform [14]. ALK is mutated or abberrently expressed in some cancers, with its ALK^ATI^ variant, characterized by an alternative transcription initiation (ATI) site, found to be expressed in 11% of melanomas [26].

Following Nellore et al, we used Snaptron to demonstrate the ALK^ATI^ variant and related EML4-ALK gene fusion can also be found in non-cancer samples. Note that whereas Nellore et al distinguish between the ALK^ATI^ variant and the EML4-ALK fusion by integrating other assays, we do not make the distinction here. We started by using Snaptron's high-level JIR query to rank samples in order according to the difference between the total coverage of ALK junctions downstream of the ATI versus the junctions upstream. The sets of upstream and downstream junctions are defined using R+F queries. We constrained the strand to be the same as that of the ALK gene and required that junctions lie within ALK’s annotated boundaries. Also following Nellore et al, we postprocessed the JIR results to exclude samples with fewer than 50 total reads covering the ALK junctions. We found that the top 10 samples in our JIR-ranked list exactly match those they reported, including the unexpected melanocyte and macrophage samples.

## 4 Discussion

Recent efforts have addressed the important and related problems of (a) analyzing many archived datasets in a uniform [13,27] and privacy-sensitive [15] fashion (b) compiling, augmenting and correcting associated metadata [28], and (c) making concise summaries available to the community [16,27]. Curated summaries are now available for collections of over 70,000 human RNA-seq samples [16]. This motivates the computational question: how do we build systems that make it easy for typical biological researchers to ask and answer questions using these resources? Snaptron is a search engine that combines summarized output from splice-aware RNA-seq alignment tools like Rail-RNA [13] with a range of indexing strategies and a sophisticated query planner. Through its array of interfaces, Snaptron allows researchers to query and visualize the vast amount of splicing data now available in summaries like `intropolis` [14]. At no point are users required to download or process raw sequencing data, or any other large files.

Snaptron’s design addresses the question of how to combine the best qualities of multiple indexing and database systems in a way that allows rapid queries, even when queries are concerned with a combination of both structured interval and numeric data, and much less structured textual metadata. The design mixes genomics-oriented software like Tabix [17] with more generic database and indexing systems like SQLite [19] and Lucene [20]. Generic systems like SQLite performed surprisingly well, sometimes better than genomics-oriented tools when queries conjoined interval constraints with other constraints.

To demonstrate its utility, we used Snaptron to assess: (a) prevalence of putative novel junctions and exons, (b) tissue specificity of novel splicing events, and (c) which public samples exhibit the most divergent splicing patterns for a particular gene. With the growing popularity of RNA-seq analysis tools that quantify with respect to a given gene annotation [29,30], thereby trusting the completeness and accuracy of the annotation, questions about which annotated and unannotated splicing patterns are well supported by public data are increasingly crucial. Snaptron makes it easy to visualize and measure support for putative splicing patterns.

In the future it will be important to further optimize Snaptron queries that impose complex constraints on both sample metadata and junction data. While we proposed and implemented an Aho-Corasick-based method that is efficient for some expected queries — with response times measured in seconds or tens of seconds — it is not hard to construct more complex queries that push response times to minutes. The main example of this kind of search type is where thousands of samples are used to further filter a larger R+M or R+F+M query where 100’s of thousands of junctions are returned. A specific example would be querying for all junctions on chromosome 1 while also requiring that all junctions appear in samples that have the keyword “tissue” in their description metadata field in the GTEx compilation. The problem is fundamentally difficult since it requires scanning, compiling and summarizing a large fraction of the overall data compilation. It will be important to investigate more sophisticated algorithms, caching schemes, and other methods for reducing response time.

Another future goal is to generalize Snaptron’s current support for coverage summaries like mean and SSC to additionally support user-defined functions. There are many possible summaries users could define or that have been proposed in previous studies, “percent spliced in” (PSI) [31] being a well known example. Allowing user-defined functions would require major changes, but would also obviate the need for Snaptron to individually support a large number of potential summaries.

Sample metadata quality is another important issue to be addressed in future work. The quality of metadata in public sequencing archives is generally poor, but recent efforts such as MetaSRA [32] provide both new methods as well as improved compilations of metadata. In the future, it will be important to integrate such efforts into Snaptron.

## Acknowledgments

We thank Princy Parsana for her advice when compiling TCGA metadata. We thank Leonardo Collado-Torres for his advice on the project and for his help testing the service. We thank Sarven Sabunciyan for suggesting splice junctions related to exonized repetitive elements for the second investigation. We thank Joyce Wilks, Elisabeth Wilks and Anand Malpani for their help in naming the project.

## Funding

CW and BL were supported by NIH/NIGMS grant 1R01GM118568 to BL. BL was also supported by National Science Foundation grant IIS-1349906.

